# A dipteran parasite regulates 20E synthesis and antibacterial activity of the host for development through inducing host nitric oxide production

**DOI:** 10.1101/2023.10.30.564679

**Authors:** Minli Dai, Zhe Jiang, Fanchi Li, Jing Wei, Bing Li

## Abstract

Parasites primarily rely on the host to survive, and have evolved diverse survival strategies. In the present study, we report a specific survival strategy of a dipteran parasite *Exorista sorbillans* (Diptera: Tachinidae), which is a potential biological control agent for agricultural pests and a pest in sericulture. We found that the expression levels of host nitric oxide synthase (NOS) and nitric oxide (NO) production were increased after *E. sorbillans* infection. Reducing NOS expression and NO production with a NOS inhibitor (L-NAME) in infected hosts significantly impeded the growth of *E. sorbillans* larvae. Moreover, the 20-hydroxyecdysone (20E) biosynthesis of infected hosts was elevated with increasing NO production, and inhibiting NOS expression also lowered 20E biosynthesis. More importantly, induced NO synthesis was required to eliminate intracellular bacterial pathogens that presumably competed for shared host resources. Inhibiting NOS expression down-regulated the transcription of antimicrobial peptide genes and increased the number of bacteria in infected hosts. Collectively, this study revealed a new perspective on the role of NO in host-parasite interactions and a novel mechanism for parasite regulation of host to support its development.

**Author summary:** Parasites are ubiquitous on earth and diverse, and have evolved various strategies to match their own specific requirements for survival within the host. In this study, we found that *Exorista sorbillans*, an endoparasite of many agricultural pests and the main insect pest in sericulture, used a novel strategy to manipulate its host NO synthesis for survival within its host *Bombyx mori*. Specifically, NOS expression and the production of NO in the host were up-regulated by *E. sorbillans* infection, which benefited larval *E. sorbillans* development. Meanwhile, increased NO production in host activated 20E synthesis of parasitized hosts and triggered the expression of AMPs against the invasion of pathogenic bacteria to avoid nutritional competence in the host. Our study provides innovative insights into the mechanisms by which a parasite manipulates host NO production for development and may help to expand our knowledge of other parasitic systems.

## Introduction

Parasitism is ubiquitous on earth and diverse, all living organisms are vulnerable to parasites [1]. Parasites rely on their hosts, from which they obtain benefits such as nutrition and habitat for their development [2]. Long co-evolution has driven parasites to display complex and exquisite strategies to overcome host immune response and manipulate host physiological environments to ensure their own development [3, 4]. For example, parasites utilize several immune evasion strategies and suppress immune responses of the host, or even exploit host immune system to maximize the success of their parasitic lifestyle [5, 6]. Although regulation of host by parasites is a widespread phenomenon, the diverse regulatory mechanisms remain to be explored.

Nitric oxide (NO) is produced by the breaking down of the sole substrate arginine mediated by the nitric oxide synthases (NOS) and involved in different physiological processes, including nervous transmission, vascular regulation, immune defense, and in the pathogenesis of several diseases [7–9]. In insects, NO is a primary killing factor that against many pathogens including viruses, bacteria, fungi, protozoan, and parasites. It keeps the insect immune response running effectively always by regulating antimicrobial peptides (AMPs) synthesis [10]. After parasitized by hymenopteran parasitoid, the NO level and AMP expression were induced in the host *Drosophila melanogaster*; and inhibiting NO production suppressed host AMP production and the ability to encapsulate parasitoid eggs [11–13]. In the silkworm, the microsporidia *Nosema pernyi* invasion triggered the activation of NO production and the up-regulation of AMPs, which significantly increased the ability of the host to defend against *Staphylococcus aureus* infection [10, 14]. The *Plasmodium* infection also up-regulated the NOS expression in *Anopheles stephensi*, and inhibiting host NOS activity obviously increased parasite reproduction [15, 16]. *Escherichia coli* invasion increased NO concentrations and AMPs transcriptions in *Ostrinia furnacalis*, and reducing NO level and AMPs induction decreased the survival rate of host [17]. Therefore, it is likely that the defense role of NO against parasites and microorganisms in insects is highly conserved.

There is also accumulating evidence that NO plays important roles in many developmental and physiological processes of insects, such as intestinal relaxation, neuronal communication, embryonic development, and hormone synthesis [18]. In *Apis mellifera*, it has been proved that NO is necessary for memory and olfactory habituation [19]. NO also functions in the acquisition of sperm motility in *Bombyx mori* [20]. In few insect species, such as *D. melanogaster* and *Manduca sexta*, NO is indispensable for insect development by regulating 20-hydroxyecdysone (20E) production [21, 22]. As initially subtly defined in *Drosophila*, NO directly binds ecdysone-induced protein E75 and activates hormone receptor 3 (HR3) mediated beta Fushi tarazu factor 1(βftz-f1) transcription activation, which stimulates the transcription of 20E biosynthetic genes and the subsequent 20E biosynthesis in prothoracic gland [23, 24]. Inhibition of NO production in *Drosophila* disrupts 20E production, leading to metabolic defects and a failure to initiate metamorphosis [22, 23]. Elevation of NO level significantly increases *HR3* and *βftz*-*f1* expression in larvae of *M. sexta*, which may enhance the capacity of the prothoracic gland to secrete ecdysone [25]. Of particular interest is how NO coordinates immunity and development in insects when suffering from pathogen infection.

The Diptera Tachinidae is an endoparasite, whose larvae (maggots) feed and develop inside their hosts [26, 27]. They attack a range of other insects in the orders of Lepidoptera, Coleoptera, Heteroptera, Hymenoptera and Orthoptera, such as *Mythimna separate*, *Helicoverpa armigera* and *Operophtera brumata* [28]. They have been involved in the programs of biological control against various insect pests [28, 29]. The tachinid parasitoids *Exorista japonica*, *Exorista sorbillans* and *Blepharipa zebina* are major larval endoparasites of *B. mori* and *Antheraea pernyi*, which severely damage the sericulture industry [30–32]. We have previously demonstrated that the larval development of *B. mori* is accelerated, and the expression of NOS in *B. mori* is up-regulated by *E. japonica* parasitism [33, 34], suggesting that NO might be involved in host development, which provides us a good system to study the role of NO in the host under the parasitoid stress. In the present study, the effects of *E. sorbillans* parasitism on host NO production and the role of host NO in *E. sorbillans* development were investigated. Meanwhile, the regulatory mechanism of NO mediated 20E biosynthesis and the antibacterial response of *B. mori* during parasitism were determined. This study reveals a novel mechanism by which a parasite stimulates host NO accumulation to benefit its development.

## Results

### Induced NO is not involved in host immunity against the parasitoid *E. sorbillans*

To assess whether host NO production is elevated and responsible for host immunity after *E. sorbillans* parasitism, the expression level of *BmNOS* and NO levels in host *B. mori* at 3 days after parasitism (dap) were measured. The results showed that the mRNA levels of *BmNOS1* and *BmNOS2* in parasitized hosts were significantly up-regulated by 7.40- and 1.92-fold, respectively (S1A Fig), and the NO levels were significantly increased by 2.42-fold in the hemolymph of parasitized hosts at 3 dap (Fig 1A). These data indicated that the NO-mediated host immunity might be activated in response to the tachinid parasitoid.

**Figure 1.**
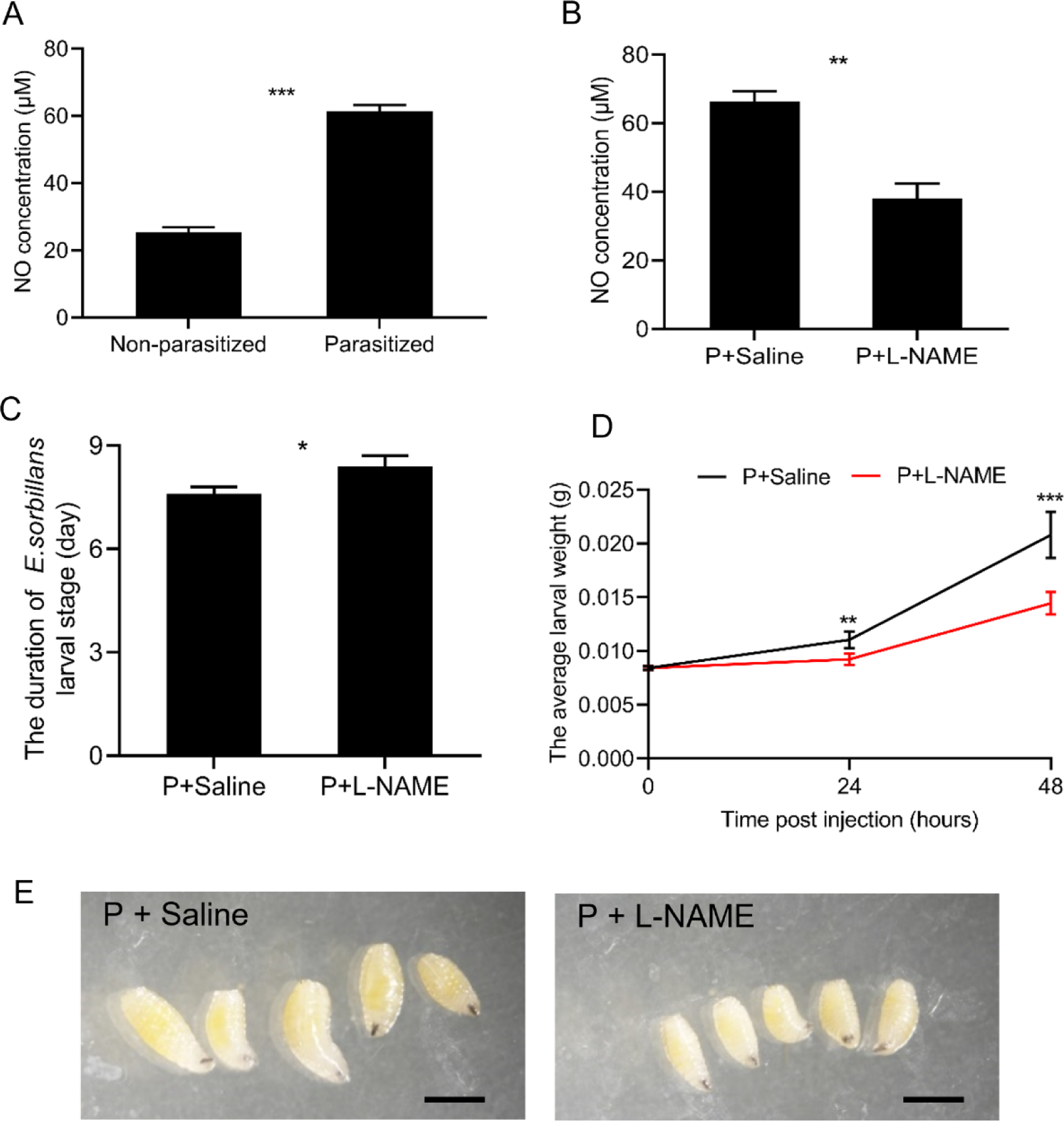
The effects of host NO on the development of *E. sorbillans* larvae. (A). Levels of NO in non-parasitized and *E. sorbillans* parasitized host larvae at 3 days after parasitism (dap). (B). NO content of parasitized (P) *B. mori* treated with saline and L-NAME at 24 h post injection. (C). The duration of the larval instar stages of *E. sorbillans* larvae in saline-treated (n = 100) and L-NAME-treated (n = 100) hosts. (D). The body weight of *E. sorbillans* larvae in saline-treated (black curve, n = 50) and L-NAME-treated (red curve, n = 50) hosts. (E). The instar of *E. sorbillans* in hosts treated with saline and L-NAME at 48 h post injection, bars, 1 mm. Data were analyzed by Student’s *t*-test and significance was determined by (\**P* < 0.05; \*\**P* < 0.01; \*\*\**P* < 0.001).

To test the above hypothesis, different concentrations of the NOS inhibitor L-NAME solution (3, 10, 30, 60 and 120 μg L-NAME/ larva) were injected into the hemocoel of hosts to reduce NOS expression and NO levels induced by parasitism. The results showed that L-NAME appeared to inhibit the transcription of *BmNOS* in a dose-dependent manner (S1B and S1C). To ensure the highest inhibitory effect, the maximum concentration of L-NAME (120 μg L-NAME/ larva) was selected to treat the parasitized hosts. The transcript levels of *BmNOS1* and *BmNOS2* in hosts were down-regulated at 24 h after L-NAME treatment (S1D). Also, the NO level was significantly lower in hosts injected with L-NAME than that in saline injected ones (Fig 1B). Intriguingly, the duration of *E*. *sorbillans* larval development in L-NAME treated hosts was extended by ∼0.79 day compared with that in the saline treated hosts (Fig 1C, *P* = 0.03). Moreover, the body weight of larval parasitoids in L-NAME treated hosts was significantly lower (Fig 1D and 1E*, P* = 0.002 for 24 h, *P* = 0.0002 for 48 h). To examine whether L-NAME had a direct inhibitory effect on larval *E*. *sorbillans* development, the larvae were reared in artificial medium supplemented with L-NAME or saline. Results showed that there was no significant difference in survival rate and body weight of larvae cultured *in vitro* under L-NAME treatment at different concentrations from 0 h to 96 h (S1E-G). These results suggested that the increased host NO production due to parasitism might benefit rather than inhibit *E*. *sorbillans* development, which was unexpected and contradictory to the typical immune function of NO in many other parasite-host systems, indicating a potential new function of NO in parasitoid-host interaction.

### *E. sorbillans* parasitism induces concurrent increase in NO production and 20E synthesis in *B. mori*

NO positively regulates 20E biosynthesis in prothoracic gland and is necessary for metamorphosis in *Drosophila* [23]. In addition, we have previously demonstrated that tachinid parasitism increased 20E titer in the host, thereby resulting in precocious onset of metamorphosis [34]. We therefore investigated whether the induced NO accounted for elevated 20E biosynthesis in *B. mori* after *E*. *sorbillans* parasitism. The RT-qPCR results showed that transcription levels of *BmNOS1* and *BmNOS2* in prothoracic gland of hosts were significantly increased by 1.83- to 5.62-fold at 1, 3 and 5 dap (Figs 2A and 2B). In accordance with up-regulated NO level at 3, 5 dap, the parasitized hosts showed a significant increase in 20E levels from 3 to 6 dap, with an almost 40% increase at 5 and 6 dap (Figs 2C and 2D). Meanwhile, the transcription levels of 20E biosynthetic genes (*BmNvd*, *BmSro*, *BmSpo*, *BmPhm*, *BmDib* and *BmShd*) in prothoracic gland were dramatically up-regulated at 1, 3 and 5 dap (Fig 2E). Next, the effect of NOS inhibitor on 20E synthesis in non-parasitized silkworms was investigated. The dose of 3 μg L-NAME did not affect the level of 20E, whereas the doses of 30, 60 and 120 μg L-NAME significantly reduced 20E titers (S2A Fig). Meanwhile, the transcription levels of *BmβFtz-f1* and 20E biosynthetic genes in *B. mori* prothoracic gland were decreased by 0.09- to 0.71-fold at 24 h post L-NAME treatment (S2B and S2C). As these manipulations are expected to alter the efficacy and timing of ecdysone production, we monitored the performance of *B. mori*. The saline-injected *B. mori* larvae entered wandering stage on day 8 of the 5th instar, while the duration of final instar of L-NAME treated *B. mori* larvae was significantly prolonged by ∼1 day (S2D Fig, *P* = 0.03). Collectively, these results showed that NO was required for 20E biosynthesis and normal metamorphosis in *B. mori*, and the patterns of change in 20E titers and NO levels in *B. mori* following parasitism were consistent, suggesting that parasitism induced NO might positively regulate 20E synthesis in *B. mori*.

**Figure 2.**
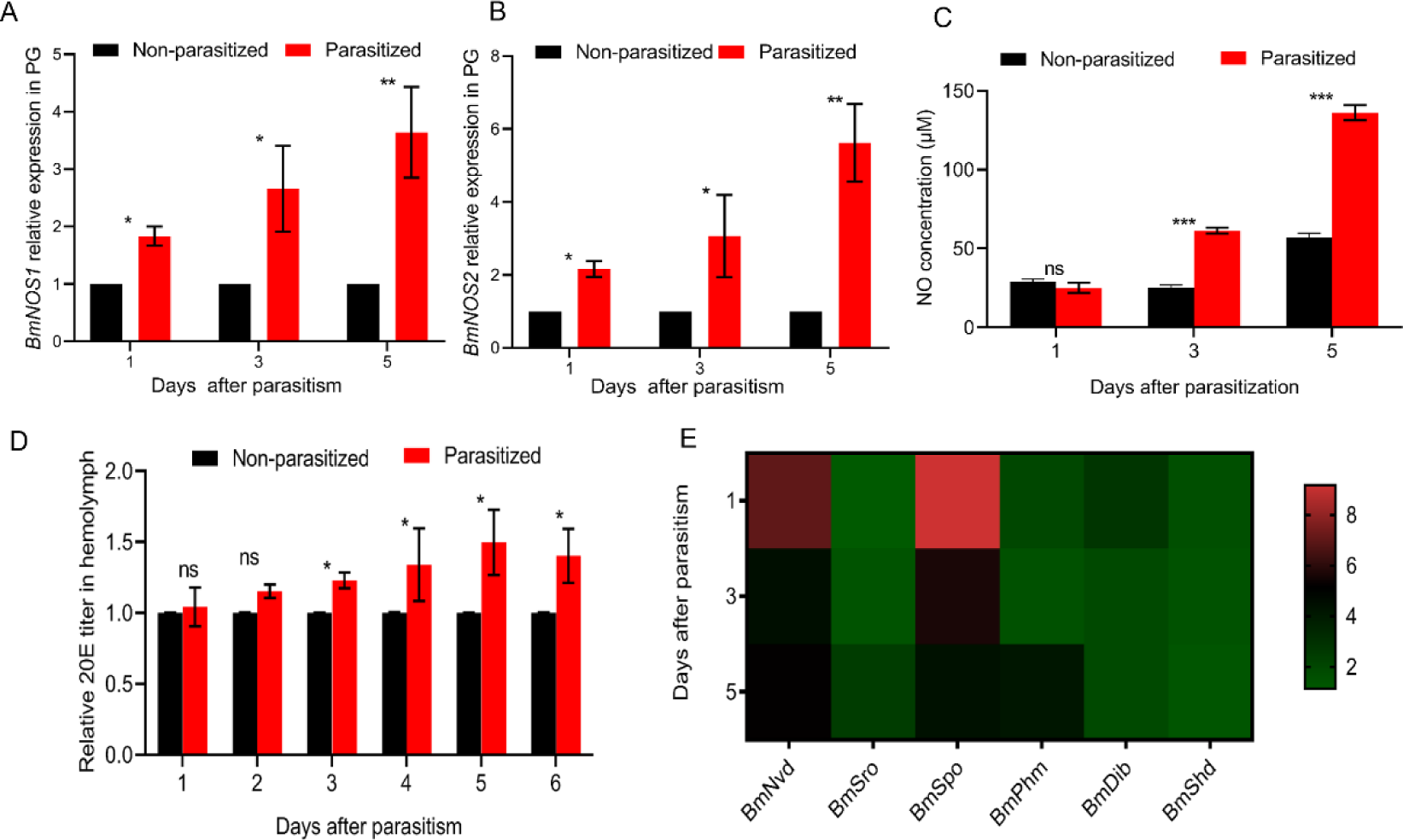
The effects of *E. sorbillans* parasitism on 20E synthesis in *B. mori*. (A, B). The expression levels of *BmNOS1* and *BmNOS2* in prothoracic gland (PG) of non-parasitized and parasitized *B. mori* at 1, 3, 5 dap. (C). NO levels in non-parasitized and parasitized host larvae at at 1, 3, 5 dap. (D). Relative levels of 20E from the hemolymph of non-parasitized and parasitized host larvae at 1 to 6 dap. (E). The transcriptional levels of 20E biosynthetic genes in PG of non-parasitized and parasitized *B. mori* at 1, 3, 5 dap. All the data were presented as mean ± SD of three independent experiments. Significance of the results was determined by the Student’s *t*-test (\**P* < 0.05; \*\**P* < 0.01; \*\*\**P* < 0.001; ns, not significant).

### Inhibition of NO production caused by parasitism impedes 20E biosynthesis in *B. mori*

To further elucidate the hypothesis that the elevated 20E level in parasitized *B. mori* was attributed to NO increase, L-NAME or insect saline was injected into host larvae at 3 dap. The results showed that the 20E titer in parasitized hosts was largely decreased by 0.75-, 0.77- and 0.78-fold at 12, 24 and 48 h post L-NAME treatment (Fig 3A). Besides, the transcription levels of *BmβFtz-f1* in prothoracic gland of hosts injected with L-NAME were down-regulated in all tested time points compared with that in hosts injected with saline (Fig 3B). Meanwhile, the expressions of 20E biosynthetic genes in L-NAME injected hosts were also dramatically suppressed (Fig 3C). Notably, concomitant with reduced 20E titer, the duration of final instar to wandering stage was significantly prolonged by ∼1 day in L-NAME treated hosts (Fig 3D, *P* = 0.02). Therefore, the positive effect of NO on 20E biosynthesis in host *B. mori* was significantly reversed by inhibition of NO production, indicating that *E. sorbillans* parasitism stimulated host NO mediated 20E biosynthesis.

**Figure 3.**
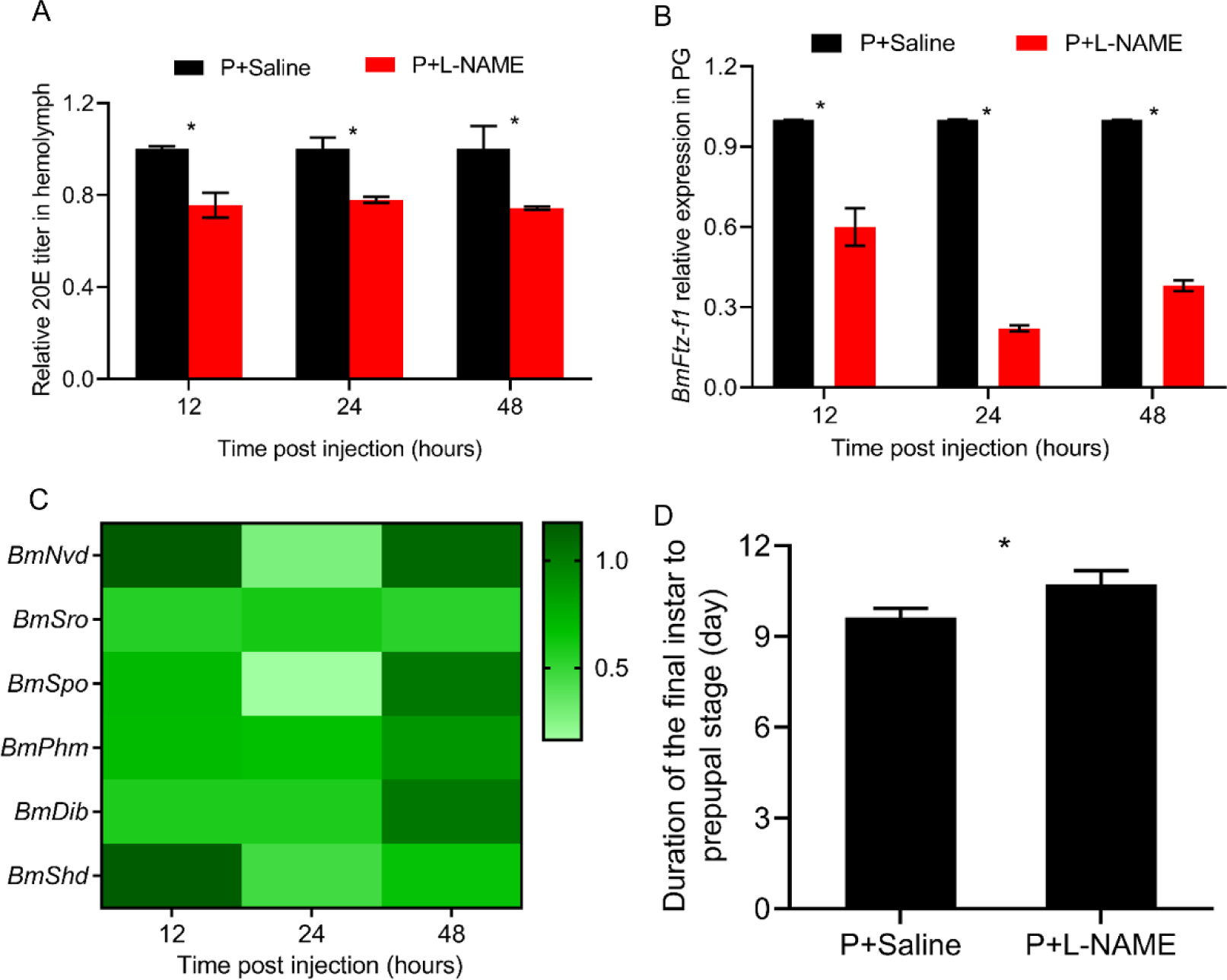
The effects of NOS inhibitor treatment on 20E synthesis in parasitized *B. mori*. (A). Measurement of the relative 20E titers in the hemolymph from parasitized host treated with saline and L-NAME at 24 h post injection at 12, 24 and 48 h post injection. (B). Relative mRNA levels of *BmβFtz-f1* in PG of the host treated with saline and L-NAME at 12, 24 and 48 h post injection. (C). The transcriptional levels of 20E biosynthetic genes in PG of parasitized *B. mori* injected with saline and L-NAME at 12, 24 and 48 h. (D). Duration of the final instar larval stage to the wandering stage in parasitized hosts treated with saline and L-NAME (120 μg L-NAME/ larva). All the data were presented as mean ± SD of three independent experiments. Significance of the results was determined by the Student’s *t*-test (\**P* < 0.05).

### Parasitism induced NO generation was involved in the antibacterial response of ***B. mori***

In a system where host and parasitoid are deeply connected, it is evident that the host might be attacked by other pathogens, like environmental bacteria that can be introduced into the hemocoel via external wound sites [35, 36]. Given the role of NO in insect immunity, we therefore deduced that parasitism induced NO elevation might also exhibit antibacterial activity in the host. To elucidate this speculation, we initially observed whether bacteria were introduced into the host hemolymph by larval parasitoid infection, the diluted hemolymph solution from the non-parasitized and parasitized *B. mori* was inoculated onto LB agar plates for bacterial proliferation examination. The results showed that extracellular bacteria had invaded the hemocoel of parasitized hosts, and the strains were identified by sequence alignment in NCBI (Figs S4A and S4B, S2 Table). Considering that the bacteria in host hemolymph might be introduced from the surface of the host body or the tachinid eggs, we sequenced the bacterial populations of these two possible originations. Results showed that both the host surface and hemolymph were colonized by the bacterial strain *Staphylococcus gallinarum strain AB328* (S1 and S2 Tables). Besides, none of the bacteria identified in freshly laid tachinid eggs were present in host hemolymph (S3 Table). RT-qPCR analysis further indicated that the expression levels of AMPs in host fat body were significantly up-regulated by 1.68- to 8.23-fold at 1 to 5 dap (Fig 4C). These results suggested that host surface bacterial populations could be introduced into host hemolymph during parasitism, and the induced expression of AMPs was probably involved in defense against the extracellular pathogenic bacteria.

**Figure 4.**
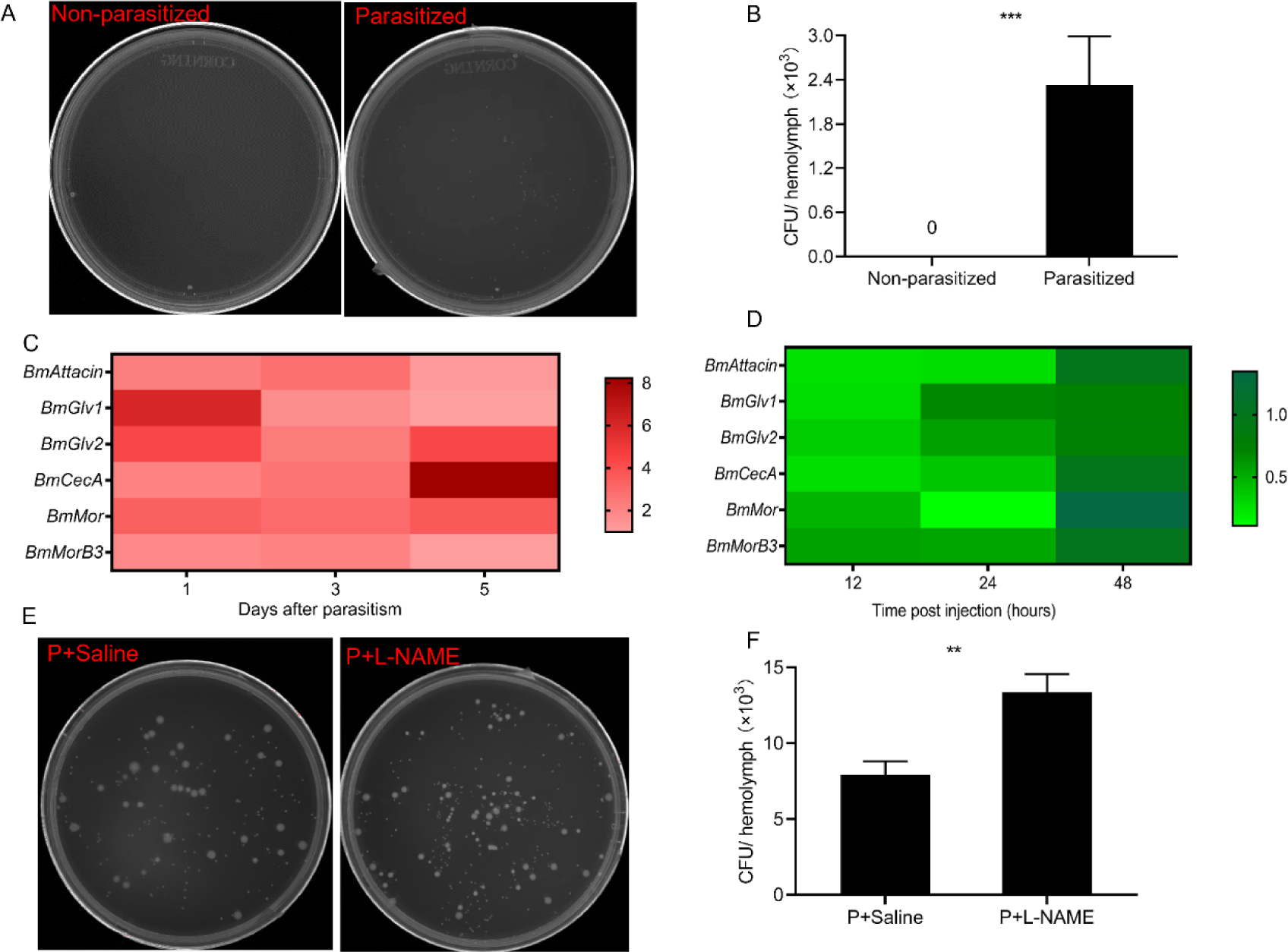
The influences of induced NO production on host antibacterial response during parasitism. (A). Bacteria in hemolymph were demonstrated by spreading non-parasitized and parasitized silkworm hemolymph on LB agar plates at 3 dap. (B). Quantification of total bacterial CFU in non-parasitized and parasitized host larvae hemolymph. (C). The transcriptional levels of AMP genes in fat body (FB) of non-parasitized and parasitized *B. mori* at 1, 3 and 5 dap. (D). Expression levels of AMP genes in FB of parasitized host injected with saline and L-NAME at 12, 24 and 48 h. (E). Bacteria from hemolymph of parasitized larvae treated with saline and L-NAME at 24 h post injection were incubated on LB agar plates. (F). Quantification of total bacterial CFU in saline- and L-NAME-treated silkworm larvae hemolymph at 24 h post injection. Significance of the results was determined by the Student’s *t*-test (\*\**P* < 0.01; \*\*\**P* < 0.001).

To further clarify whether the induced NO was involved in the increased antibacterial activity of *B. mori* after parasitism, L-NAME was injected into the parasitized hosts. RT-qPCR analysis indicated that the transcript levels of AMP genes in hosts injected with L-NAME were dramatically reduced at 12 and 24 h post injection (Fig 4D). The number of bacteria was significantly higher in hemolymph of L-NAME-injected hosts (CFU=13.37 × 10^3^) than that in saline-injected hosts (CFU=7.91 × 10^3^) at 24 h post treatment (Figs 4E and 4F). These findings suggested that parasitism induced NO production was required for the up-regulation of AMPs, which helped the host resist invading bacterial pathogens.

## Discussion

It is generally accepted that NO possesses antiparasitic effects and limits the development of parasites in the insect hosts [37, 38]. *Anopheles* mosquitoes responded to *Plasmodium* infection by up-regulating *AsNOS* expression and synthesizing NO, which limited parasite development in the midgut [39, 40]. The number of parasite *Trypanosoma cruzi* was reduced in host *Rhodnius prolixus* (Triatominae: Reduviidae) treated with NO substrate arginine, and *R. prolixus* treated with L-NAME benefited the infection and development of *T. cruzi* [41]. In addition, the level of NO in host *Drosophila* exhibited a dramatic increase immediately upon parasitoid infection, inhibiting the synthesis of NO in the host significantly increased the survival rate of hymenopteran parasitoids *Leptopilina heterotoma* and *Asobara tabida* [11, 12]. In current study, an increase in NO production was also found in the host *B. mori* infected by the dipteran parasitoid *E*. *sorbillans*, which benefited parasitoid development (Fig1 and FigS1). Likewise, as evidenced in few species, host NO acts as a friend for parasites. For example, NOS knockout rats showed a completive resistance to *Toxoplasma gondii* infection, and *Toxoplasma* was found to utilize host NO to benefit its own development and growth in the human host [42, 43]. Therefore, NO plays an inconsistent role in different host-parasite systems.

Insect parasitoids frequently modulate host development through juvenile hormone (JH) and ecdysteroid hormone pathways in order to create an environment to benefit their growth [44]. For example, *Cardiochiles nigriceps* (Hymenoptera: Braconidae) parasitism caused delayed and impaired pupation by reducing 20E titer in host *Heliothis virescens* (Lepidoptera: Noctuidae) [45]. In contrast, a few parasitoid species could induce a shorten development duration or a precocious metamorphosis through increasing the 20E level in host or decrease host JH concentration, such as *Chaetophthalmus dorsalis* (Diptera: Tachinidae), *Microplitis pallidipes* (Hymenoptera: Braconidae), *Chelonus inanitusid* (Hymenoptera: Braconidae), etc [46–48]. In this study, consistent with *E. japonica* infection, *E. sorbillans* parasitism elevated 20E titers in the host *B. mori*, indicating that changes in host 20E level in response to parasitism were a broad event in the dipteran *Exorista* species [34]. Insect 20E synthesis was regulated by multiple signaling pathways such as MAPK signaling pathway, activin signaling pathway, PI3K/AKT pathway, NO signaling pathway, TOR signaling, etc [49]. Our previous transcriptome analysis showed that differentially expressed genes were significantly enriched in NO signaling pathway in *B. mori* after *E. japonica* parasitism [33]. We also observed an increase in *BmNOS* expression levels and NO production in *B. mori* following *E. sorbillans* parasitism, which accounted for the increased host 20E titer (Fig 2). Additionally, miRNAs produced by the bracovirus of parasitoid wasp *Cotesia vestalis* could directly alter 20E signal and inhibit the growth of the host [50]. Here, we found that tachinid larvae manipulated host 20E synthesis and development to benefit their growth by indirectly inducing host NO production.

The host infected by one pathogen is more likely to suffer the infection of contemporary pathogens, and both pathogens utilize the nutrients in the host, leading to competition of host resource between these pathogens [6, 51]. In host *Helicoverpa armigera* (Lepidoptera: Noctuidae), the entomopathogenic fungus *Metarhizium rileyi* infection induced the invasion of opportunistic bacteria into host hemocoel, leading to competition between *M. rileyi* and bacteria for amino acids in the host hemolymph. Meanwhile, *M. rileyi* activated the AMPs expression and exploited antibacterial activity of the host to eliminate bacteria competing for nutrients [6]. *Polistes dominulus* wasps (Hymenoptera: Vespidae) were more efficient at eliminating bacteria when they were parasitized by the endoparasite *Xenos vesparum* (Strepsiptera: Xenidae), which caused by the up-regulation of host defensin after parasitism [35]. In present study, *E. sorbillans* infection introduced environmental bacterial populations into the host and the bacteria presumably competed for host resources. The up-regulation of AMPs mediated by parasitism induced NO production inhibited the growth of pathogenic bacteria (Fig 4). Indeed, it is a fundamental feature for parasites to exploit nutrition of host insects. From the evolutionary perspective, restricting the growth of pathogenic bacteria can avoid nutrient competition and thus facilitate the development of parasites.

The evolutionary consequence of a parasitic life style makes the parasites have the ability to maximally exploit host resources and minimize their own energy inputs [52]. During the co-evolution process, it has been proved in several parasitoid insects that the ability to biosynthesize nutrients was lost. For example, several parasitoid species such as *Nasonia vitripennis* were lack of lipogenesis, instead, they stimulated host lipid accumulation that could be subsequently utilized by parasitoids [53, 54]. The wasp *Cotesia chilonis* could not synthesize ten essential amino acids because of the absence of key genes in the biosynthetic pathway, in turn, the wasp induced the increase in amino acid content in host hemolymph to satisfy its own amino acid requirement [55]. *In vitro* rearing of *E. sorbillans* showed that larval-pupal-adult transition only occurred with the supplementation of exogeneous 20E, which implied that they were not able to synthesize sufficient 20E to grow and initiate metamorphosis [56]. In present study, *E. sorbillans* parasitism enhanced host 20E synthesis through inducing NO production, inhibiting host NO production decreased 20E titer, suggesting that normal progress of larval growth relied on host 20E that stimulated by NO (Fig 3). Our previous study also demonstrated that *E. japonica* parasitism could elevate host 20E synthesis [34]. Therefore, we deduce that the tachinid parasitoids probably lack the elements involved in 20E synthesis, which can be further determined by genomic sequencing of these parasitoids. In addition, our results also raise a question as to why parasitoids accelerate host development to compete for resources. Indeed, the death of hosts parasitized by *E. sorbillans* occurred during spinning stage to pupal stage, which restricted resource investment on host metamorphosis [57]. Thus, we presume that the adaptive strategy for tachinid parasitoid is to stimulate host 20E accumulation for its larval development, which although temporarily promotes host development, the growing parasitoid larva consumes a large amount of host nutrients and causes host death rapidly after exploitation of host 20E. Our results bring new insights into a novel mechanism for tachinid parasite manipulation of host 20E biosynthesis by inducing the NO production to promote development and parasitic efficiency.

In conclusion, our results highlight the critical function of NO in parasitoid development. We found that *E. sorbillans* parasitism activated NOS and the production of NO. Increased NO production in host conferred no resistance to *E. sorbillans* development, but activated 20E synthesis of parasitized hosts and triggered the expression of AMPs against the invasion of pathogenic bacteria to avoid nutritional competence in the host (Fig 5). Our study adds to the existing knowledge of NO function in development and immunity of insects, and also sheds new light on the host-parasite association.

**Figure 5.**
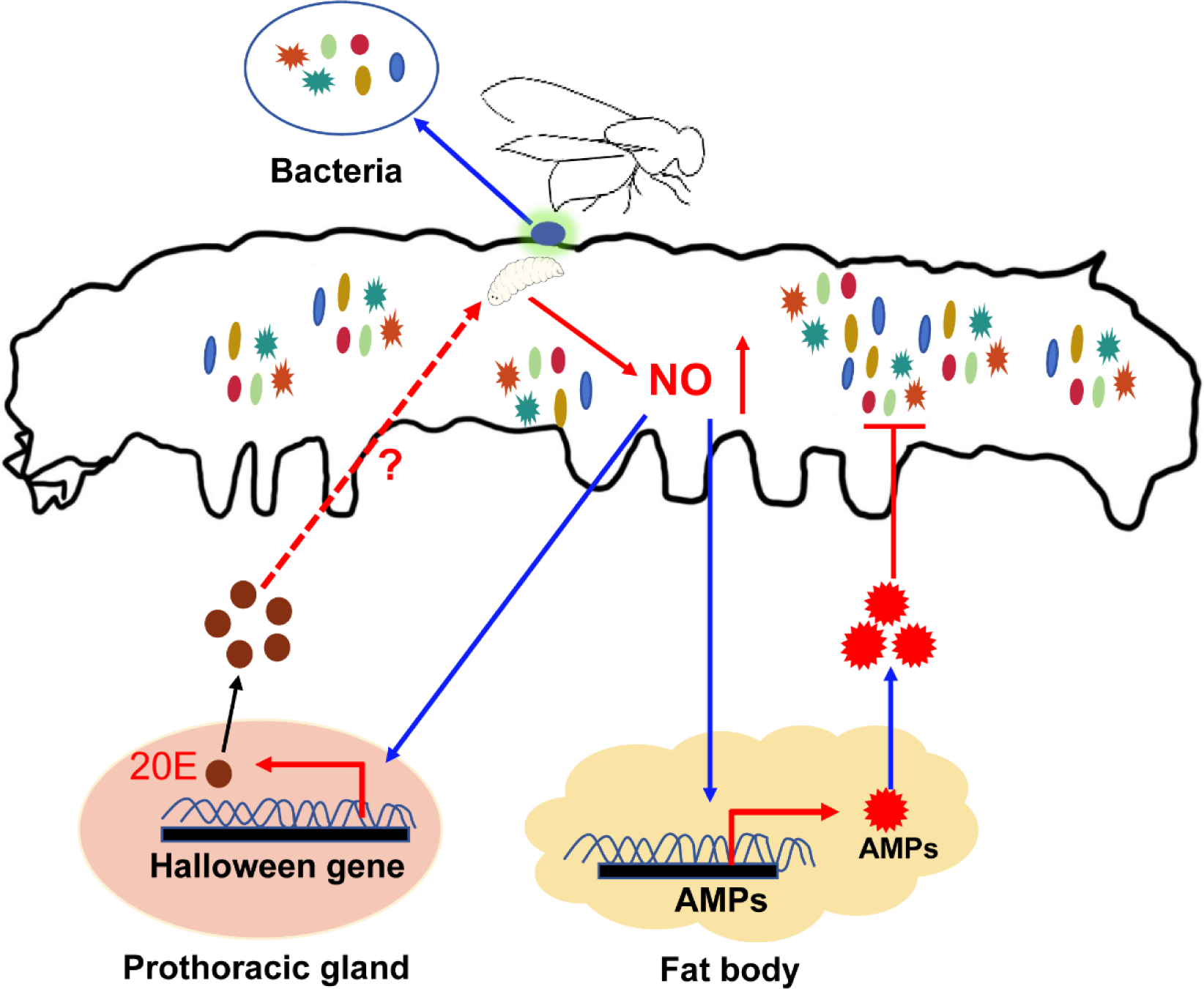
A schematic model of dipteran parasitoid *E. sorbillans* manipulates 20E synthesis and antibacterial activity of the host *B. mori* for development through inducing NO production. The *NOS* transcription and NO level in host *B. mori* were significantly increased after *E. sorbillans* parasitism, whereas, the increased NO production in host possesses no resistance to *E. sorbillans* larvae but activates 20E synthesis of parasitized host. Meanwhile, the elevated-NO level triggers the expression of AMPs, which might function in the elimination of pathogenic bacteria to avoid nutritional competence in host. The efficient anti-bacterial response therefore prevents bacteria from competing with *E. sorbillans* for the host nutrients required for its own growth.

## Materials and methods

### Insects rearing and tissue collection

The silkworm *B. mori* (Jingsong × Haoyue strain) and tachinid fly *E. sorbillans* used in this study were stocked in our laboratory and reared as previously described [34]. Briefly, the *B. mori* was reared on mulberry leaves at 25 ± 1 °C, 75 ± 5% relative humidity, and a 12 hour light:12 hour dark photoperiod. The *E. sorbillans* colony was maintained in our laboratory by allowing mated female flies to parasitize the fifth instar silkworms. The mature larvae were finally egressed from the silkworms and pupated, and the pupae were placed in a net cage (45 × 45 × 45 cm). The emerging adults were fed with 10% honey solution. After mating, each of the cages was loaded with 50 silkworm larvae on day 1 at the fifth instar for parasitism and the parasitized hosts were collected for the subsequent experiments.

The fat body, hemolymph, and prothoracic gland samples were collected from silkworm individuals at 1, 3 and 5 days after parasitism (dap) and the non-parasitized controls. Ten silkworm tissues (male: female =1:1) were pooled as one biological replicate three replicates for each trial were prepared and stored at -80°C for further analysis.

### Treatment of silkworms with the NOS inhibitor

The pharmacologic NOS inhibitor NG-nitro-L-arginine methyl ester hydrochloride (L-NAME) was purchased from MCE (MedChemExpress, USA) and dissolved in insect saline [10]. The concentration of L-NAME used in this study was made a reference to previous study, the silkworms were injected with different doses of L-NAME (3, 10, 30, 60 and 120 μg L-NAME/ larva) and saline treatment was applied as the control [10]. To test the effect of host NO production on the development of parasitoid larvae, the host larvae were microinjected with L-NAME solution (120 μg/ larva) or insect saline (control) at 3 dap. After injection, the hosts were reared in a 25 °C incubator until the *E. sorbillans* pupa emerged. The body weight of *E. sorbillans* larvae and the larval period were recorded. Meanwhile, the fat body and hemolymph from L-NAME-injected and saline-injected *B. mori* larvae were collected at 12, 24 and 48 hours (h) post injection.

To determine the role of NO production on 20E synthesis of *B. mori*, the prothoracic gland and hemolymph samples of larvae from the L-NAME and saline treated silkworms were collected at 24 h after injection. The development course of silkworms was also recorded.

### *In vitro* rearing of *E. sorbillans* larvae with L-NAME

To verify the direct effects of L-NAME on larval *E. sorbillans* development, the 2nd instar *E. sorbillans* was cultured with the artificial medium based on the protocols in previous report [56]. The Grace’s Insect Medium (Sigma, USA) fortified with the 10% *B. mori* hemolymph and 10% fetal bovine serum was used as the medium (500 μL in total). Different concentrations of L-NAME (0, 0.3, 3, 6, 12 and 24 μg/ μL) were applied to assess the effects of NO on parasitoid development [58, 59]. Silkworms were surfaced-sterilized with 75% ethanol and dissected at 3 dap to obtain the 2nd instar maggots. The maggots were washed 5 times with phosphate-buffered saline (PBS) and then transferred to the 6-well tissue culture plate containing the mixed rearing medium. Ten maggots were pooled as one biological replicate and five replicates were performed. All plates were incubated at 25 ± 1 °C for determining the parasitoid performance. The survival rate and body weight of *E. sorbillans* larvae were recorded. The experiments were performed three times.

### Determination of NO production

The NO level in *B. mori* hemolymph samples were measured with the Nitrate/Nitrite Assay Kit (Beyotime Biotechnology, China). A volume of 200 μL of hemolymph was collected from each *B. mori* larva. Hemolymph samples from five individuals were pooled as one tested sample and centrifuged at 12,000 rpm for 10 min at 4°C. The supernatants (60 μL) were collected for measuring NO concentrations. According to the instructions of the manufacturer, the nitrites at final concentrations of 0, 2, 5, 10, 20, 40, 60, and 80 μM in a 200 μL reaction volume were prepared for a standard curve to quantify nitrite concentrations in the samples. The absorbance at 540 nm on a microplate reader (Thermo Fisher Scientific, USA) was determined. This experiment was performed in triplicate.

### 20-hydroxyecdysone (20E) content measurement

The sample for 20E measurement was prepared according to the previously described method [34]. One hemolymph sample (500 μL each) was collected from five *B. mori* larvae by cutting an abdominal leg on ice. A total of 500 μL of hemolymph was diluted with 1.5 mL chilled methanol, and then centrifuged at 12,000 × g for 10 min [6]. The supernatants were transferred into a new tube, followed by drying with an evaporator and resuspended in enzyme immunoassay (EIA) buffer. The level of 20E in the hemolymph was determined by a commercial 20E enzyme immunoassay kit (Lvye Biotechnology, China) following the manufacturer’s instructions. The absorbance at 450 nm was determined, and the experiments were repeated three times.

### Detection of hemolymph bacteria

Non-parasitized and parasitized host larvae were randomly selected, and their hemolymph samples were collected at 3 dap. The larval *B. mori* was surface-sterilized with 75% ethanol and then rinsed three times with PBS. Freshly hemolymph samples from five larvae were collected and pooled as one sample, three samples were prepared. Each sample was diluted 100-fold in double distilled H_2_O (ddH_2_O). Subsequently, 40 μL of each dilution was spread onto LB agar plates and then incubated at 37 °C overnight [60]. The number of bacteria on each plate was recorded and the colony-forming unit (CFU) method was used to determine the total number of hemolymph bacteria. To obtain the bacteria on the surface of *B. mori*, 1 mL ddH_2_O was used to wash the silkworms, followed by centrifuged at 4,500 × g for 5 min. The supernatants were removed, then the bacterial precipitates were resuspended with 500 μL ddH_2_O and cultured on LB agar plates at 37 °C overnight [60, 61]. In order to collect the bacteria of *E. sorbillans* eggs, the silkworm was surface-sterilized with 75% ethanol and deposited in the fly cage to receive the freshly laid eggs. The eggs were collected, surface-sterilized with 75% ethanol, washed with ddH_2_O, homogenized with 500 μL ddH_2_O and plated onto LB agar plates, incubated at 37 °C overnight [60, 61]. The colonies were selected randomly and streaked for purification. Genomic DNA of the different bacterial isolates was extracted with the TIANamp Bactreia DNA kit (Tiangen, China) by following the manufacturer’s instructions and used to amplify 16S rRNA gene sequences by PCR with the universal 16S rRNA specific primers (27F: AGAGTTTGATCCTGGCTCAG, 1492R: AGAGTTTGATCCTGGCTCAG) [61]. The purified PCR products were sequenced (Sangon Biotech, Shanghai), and results were aligned via BLAST searches in the National Center for Biotechnology Information (NCBI, http://blast.ncbi.nlm.nih.gov/).

### Expression analysis by reverse transcription-quantitative PCR (RT-qPCR)

The total RNA was extracted from the silkworm tissues using TRIzol reagent (TaKaRa, Japan) to analyze expression levels of *BmNOS1, BmNOS2*, 20E biosynthetic genes (*BmNvd, BmSro, BmSpo, BmPhm, BmDib* and *BmShd*), *BmβFtz-f1* and AMP genes (*BmAtta*, *BmGlv1*, *BmGlv2*, *BmCecA*, *BmMor*, and *BmMorB3*). The cDNA was prepared from the extracted RNAs with oligo (dT) and random primers using the PrimeScript RT reagent kit (TaKaRa, Japan) according to the manufacturer’s instructions. The specific primers for each gene were listed in S1 Table. RT-qPCR reactions were conducted on the Viia 7 Real-time PCR System (Applied Biosystems, USA) with SYBR Premix Ex TaqTM II (TaKaRa, Japan). The RT-qPCR was performed for at least three biological replicates under the following conditions: 45 cycles of 95 °C for 5 s, 60 °C for 10 s and 72 °C for 10 s. The expression levels of target genes were normalized with the reference gene *BmRp49* and analyzed using the 2^-ΔΔCt^ method [62].

## Statistical analysis

All statistical analyses were performed using SPSS 16.0 software (SPSS, Chicago, IL, USA). The significant difference was determined by one-way analysis of variance (ANOVA) coupled with a Tukey’ test. The *P*-value of < 0.05 *, 0.01 ** or 0.001 *** was considered significant, highly or the most highly significant, respectively. All figures were drawn by using GraphPad Prism version 7 (San Diego, USA).

## Data accessibility

The authors declare that all data supporting the findings of this study are available in the manuscript and its supplementary data files or are available from the corresponding author upon request.

## Authors’ contributions

M.D.: conceptualization, data curation, formal analysis, investigation, methodology, writing—original draft, writing—review and editing; Z.J.: data curation, formal analysis, investigation, methodology, writing—original draft; F.L.: investigation, methodology; W.J.: conceptualization, project administration, supervision, funding acquisition, writing—review and editing; B.L.: conceptualization, project administration, funding acquisition, supervision, writing—review and editing.

## Acknowledgements

We thank Zhongbin Qian from Rugao sericultural station for providing the *E. sorbillans* colony.

## Supplementary material

**Supplementary figure 1. The effects of induced host NO on the development of *E. sorbillans* larvae.** (A). The expression levels of *BmNOS1* and *BmNOS2* in FB of non-parasitized *B. mori* and parasitized ones at 3 dap. (B, C). The mRNA levels of *BmNOS1* and *BmNOS2* in FB of non-parasitized *B. mori* injected with different concentrations of NOS inhibitor at 24 h post injection. (D). Expression levels of *BmNOS1* and *BmNOS2* in FB of parasitized *B. mori* injected with NOS inhibitor (120 μg L-NAME/ larva) at 24 h post injection. (E). *In vitro* rearing of *E. sorbillans* larvae. (F). Survival rates of *E. sorbillans* larvae developed on rearing media with different doses of NOS inhibitor (n = 30). (G). The body weight changes of *E. sorbillans* larvae developed on rearing media with different doses of NOS inhibitor (n = 30). Student’s *t*-test was used for two-group comparisons and significance was determined by (\**P* < 0.05; \*\**P* < 0.01, \*\*\**P* < 0.001). Data comprising more than two groups were analyzed using ANOVA coupled with Tukey’s multiple comparison test.

**Supplementary figure 2. The influences of NOS inhibiter on 20E synthesis in *B. mori*.** (A). Relative titers of 20E from the hemolymph of non-parasitized *B. mori* treated with different concentrations of NOS inhibitor at 24 h post injection. (B). The expression level of *BmβFtz-f1* in PG of non-parasitized silkworm treated with different concentrations of NOS inhibitor at 24 h post injection. (C). Relative mRNA levels of 20E biosynthetic genes in PG of non-parasitized *B. mori* treated with NOS inhibitor (120 μg L-NAME/ larva) at 24 h post injection. (D). Duration of the final instar larval stage to the wandering stage in non-parasitized *B. mori* injected with saline and L-NAME (120 μg L-NAME/ larva). All the data were presented as mean ± SD of three independent experiments. Significance for two-group comparisons was determined by the Student’s *t*-test (\**P* < 0.05; \*\**P* < 0.01; ns, not significant), and data comprising more than two groups were analyzed using ANOVA coupled with Tukey’s multiple comparison test.

